# Metagenomic analyses of the plastisphere reveals a common functional potential across oceans

**DOI:** 10.1101/2024.08.29.610283

**Authors:** Stefan Lips, Mechthild Schmitt-Jansen, Erik Borchert

## Abstract

Trillions of plastic particles have accumulated in the oceans, covered by microbial biofilms (termed the “plastisphere”) whose functional potential remains unexplored. We evaluated genome-resolved bacterial metagenomes of the plastisphere from the North Atlantic and North Pacific garbage patches and compared their structure and functional potential to ambient plankton. Our data revealed a characteristic genetic potential of the plastisphere with functionally equivalent taxonomic units across both oceans. We found fewer coding genes, smaller genome sizes and lower GC-content in plankton in comparison to the plastisphere, despite residing in the same environment, reflecting a greater nutrient demand in the plastisphere. A functional gene analysis confirmed that the plastisphere consists of microorganisms with a higher potential to control their nutrient supply, metabolize a wider range of carbon sources, attenuate free radicals and use alternative energy sources like anoxygenic photosynthesis. Our results suggest that the overriding factor for the high functional similarity of the plastisphere in both oceans is the habitat for biofilm formation with the potential to support mutualism and nutrient sharing making genomic streamlining as found in plankton, unnecessary. Consequently, increasing plastic pollution promotes the expansion of new functional units within oligotrophic oceans, with the potential to impact biogeochemical cycles.

## Introduction

Trillions of plastic particles have accumulated in the oceans, especially in the major ocean gyres^1,2^, raising the question if marine plastic pollution is a global boundary threat^3^. So far, the risk of physical effects of drifting plastics due to ingestion, entanglement^4^ and potential hazards due to the release of additives^5,6^ or adsorbed environmental chemicals^7^ have already been identified, however a disruptive effect on ecosystem functioning remains to be demonstrated. Recognizing plastic pollution as a global boundary threat would be a first step towards the sustainable use of the oceans, seas and marine resources in the interests of sustainable development (SDG 14, https://www.refworld.org/legal/resolution/unga/2015/en/111816).

The ever-increasing amount of artificial plastic surfaces has introduced a novel entity^8,9^, especially in offshore regions where natural substrates are scarce. These surfaces are heavily overgrown by microbial biofilms, which impacts fate and potential effects of plastic^10^. This biofilm is collectively termed the ‘plastisphere’^11^ or ‘neopelagic community’^12^, as it differs from the planktonic community in species composition by 16S rRNA gene analysis^13–16^. Plastisphere biomass was estimated to range from 1000 to 15000 tons, corresponding to a total number of 2.1 × 10^21^ to 3.4 × 10^21^ cells^17^. Although this may represent ∼ 1% of the global ocean neuston surface microlayer, little is known about the plastisphere microbial ecology to successfully colonize plastic surfaces in oligotrophic oceans. Knowledge of the functional genetic potential and selected adaptation mechanisms of these communities is crucial for understanding the impact on global ecosystem functioning, such as elemental cycling.

The subtropical gyres are large systems of rotating ocean currents, in which water is transported downwards, carrying nutrients away from the photic zone and trapping the floating plastic on the surface^18^. The resulting oligotrophic areas have been designated “ocean deserts” due to their excess of solar energy and depletion of inorganic nutrients^19,20^ as well as organic carbon. Planktonic species living in the gyres have adapted to the limiting conditions by I) reducing their genome sizes and cell volume (genomic streamlining), II) replacing or minimizing the use of limiting elements (such as nitrogen, iron, phosphorus) or III) acquiring alternative metabolic pathways (such as photoheterotrophy or nitrogen fixation)^21^. These adaptations in plankton are the result of long-standing evolutionary processes, however, it is not known whether the plastisphere harbors organisms with similar traits due to a lack of functional studies. In contrast, increased amounts of chlorophyll *a* were detected on plastic in the North Pacific garbage patch^22^, suggesting significant autotrophic biomass and organic carbon production forming a eutrophic niche in an oligotrophic system. By floating on the sea surface for years before fragmentation and eventual sinking^23,24^, plastic and the adhering plastisphere are constantly exposed to high light and UV-irradiation, which can cause strand breaks in the polymer-backbone as well as biological damage such as DNA strand breakage and the formation of free radicals such as reactive oxygen (ROS) or nitrogen species (RNS). This promotes the leaching of dissolved organic carbon molecules from plastic^25^, which can be metabolized by microorganisms^26^, rendering plastic a potential source of organic carbon.

Thus, we hypothesize that the plastisphere selects for microorganisms with a different functional potential to efficiently exploit this new ecological niche in an otherwise oligotrophic environment. To test this hypothesis, we compared the functional potential based on genome-resolved metagenomes of both the plastisphere and planktonic bacterial communities of the open oceans. This includes data from geographically distinct plastisphere microbial communities from the North Atlantic garbage patch (NAGP), the North Pacific garbage patch (NPGP) sampled in summer 2019 as well as the surrounding plankton obtained from the TARA Oceans Project^27^. We specifically address the questions whether I) the plastisphere has a taxonomic and functional core composition across oceans; II) the genome of the plastisphere is genetically streamlined analogous to the plankton community; and III) the metagenomic potential of the plastisphere allows it to compensate for the oligotrophic conditions of the gyres.

Here we use a comparative metagenomic approach that examines the taxonomic and genetic differences between the plastisphere and the regional planktonic community at different geographical sites in the NAGP and NPGP (Fig. 1a). We make use of metagenome-assembled genomes (MAG) to investigate the abundance of genes from central metabolism as well as genes favoring the survival under extreme oligotrophic conditions in the gyres. We especially focus on the supply of macro- and micro-nutrients (nitrogen, phosphorus, carbon and essential elements) as well as mitigation of high-light conditions. We find systematic differences in taxonomy, genome structure and functional gene abundance between plastisphere and plankton, which suggest that the plastisphere is a new functional unit.

**Fig. 1.**
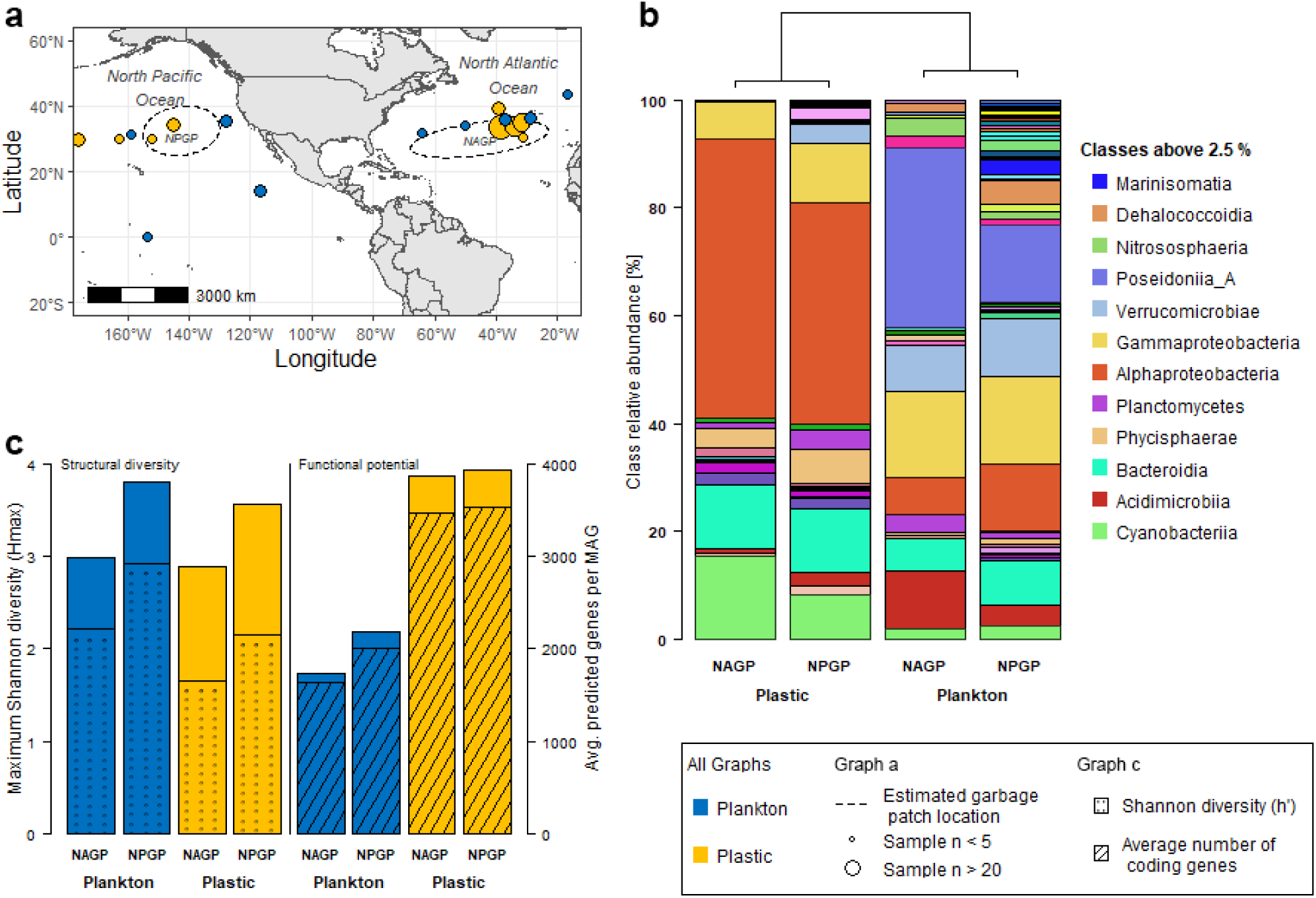
Study area and the taxonomic composition. **a** Map of the sampling sites and the modelled center of the North Pacific (NPGP) and North Atlantic garbage patch (NAPG)^28^. **b** Taxonomic comparison of bacterial classes with at least 2.5% relative abundance at the different sampling sites. Hierarchical clustering by Euclidean distance and the minimum variance method (Ward.2) distinguished plastic and planktonic habitats. **c** Comparative bar plot of plankton and plastic community diversity for structure (Shannon diversity h and maximum Shannon diversity H_max_) and the diversity of the functional potential, presented as average number of predicted genes per MAG.

## Results

### Taxonomic comparison of Atlantic and Pacific MAGs

The MAGs of plastic and plankton were sourced from individual metagenomic samples and treated equally. The Atlantic data set comprised 736 plastisphere and 213 planktonic MAGs, the Pacific data set consisted of 1021 plastisphere and 289 planktonic MAGs. Genome-to-genome comparison by FastANI analysis suggests a high degree of genome similarity between the individual samples implicating equivalent functional traits across oceans (Supplementary Table 1).

The plastisphere and plankton MAGs were composed of 29 unique phyla, which resulted from 11 phyla on NAGP plastic, 16 phyla on NPGP plastic, 17 phyla in NAGP plankton and 26 phyla in NPGP plankton samples. At the taxonomic levels from phylum to family, each habitat formed a distinct cluster with dominance of the *Alphaproteobacteria* class, which contributed 41 to 52% of the MAGs in the plastisphere and 7 to 12% in plankton (Supplementary Table 2, Fig. 1b). *Alphaproteobacteria* were dominated by the *Rhodobacteraceae* family, which accounted for 21 to 25% of MAGs in the plastisphere. The second biggest class in the plastisphere were *Bacteroidia,* accounting for 12%, followed by *Gammaproteobacteria* and *Cyanobacteria*, ranging from 7 to 15%. The *Bacteroidia* of the plastisphere were dominated by *Saprospiraceae*, *Cyclobacteriaceae* and *Flavobacteriaceae* and *Gammaproteobacteria* by *Vibrionaceae*, *Woeseiaceae*, *Alteromonadaceae,* and in the NPGP also by *Halieaceae*. The most abundant families in *Cyanobacteria* were *Nostocaceae*, *Xenococcaceae* and *Phormidesmiaceae*.

In contrast, *Gammaproteobacteria*, *Alphaproteobacteria* and *Bacteroidia* were similarly abundant in plankton with 6 to 16%. *Cyanobacteria* were very poorly represented in plankton, accounting for 1.9% in NAGP and 2.4% in NPGP. In general, plankton showed a very diverse composition at the class level, and only *Poseidonia*, which belong to the Archaea, were represented more frequently, up to 33%. Archaeal MAGs were found exclusively in plankton. Thus, the Shannon diversity index (H’) based on class data was on average 0.7 higher in plankton than in the plastisphere (Fig. 1c). In addition, species were more homogeneously distributed within the classes, resulting in a species evenness of 0.75 in plankton, 0.17 higher than in the plastisphere. The genetic composition of the communities showed large differences, with the planktonic species having on average 50% fewer predicted genes per MAG than the plastisphere. Moreover, the proportion of coding genes was about 3% higher in the planktonic species. Considering the greater number of predicted genes per MAG in the plastisphere, this coding difference results in three times more non-coding genes in plastic MAGs.

### Comparison of genome size and GC-content

The plastisphere had a significantly higher median genome size (p < 10^-20^) of 3.4 ± 1.4 Mb, compared to 1.8 ± 0.9 Mb in planktonic MAGs (Fig. 2a). Moreover, the GC-content was significantly higher in the plastisphere MAGs (p < 10^-20^), with a median of 58.4 ± 8.1% compared to 44.7 ± 9.0% in plankton (Fig. 2b). ANOVA analysis revealed only a slight impact of the respective ocean basin on the genome size (p < 10^-3^) and no impact on GC-content. Even for taxonomically identical classes, the species on plastic exhibited a 2.3-fold larger genome and a 1.3-fold higher GC-content (Supplementary Table 3). This was mostly due to a divergent taxonomical composition in the lower taxonomic levels. Only the *Bacteroidia* and *Alphaproteobacteria* were dominated in both habitats by similar families such as *Flavobacteriaceae* or *Rhodobacteraceae*, but their genera also differed and had a lower GC-content and smaller genomes.

**Fig. 2.**
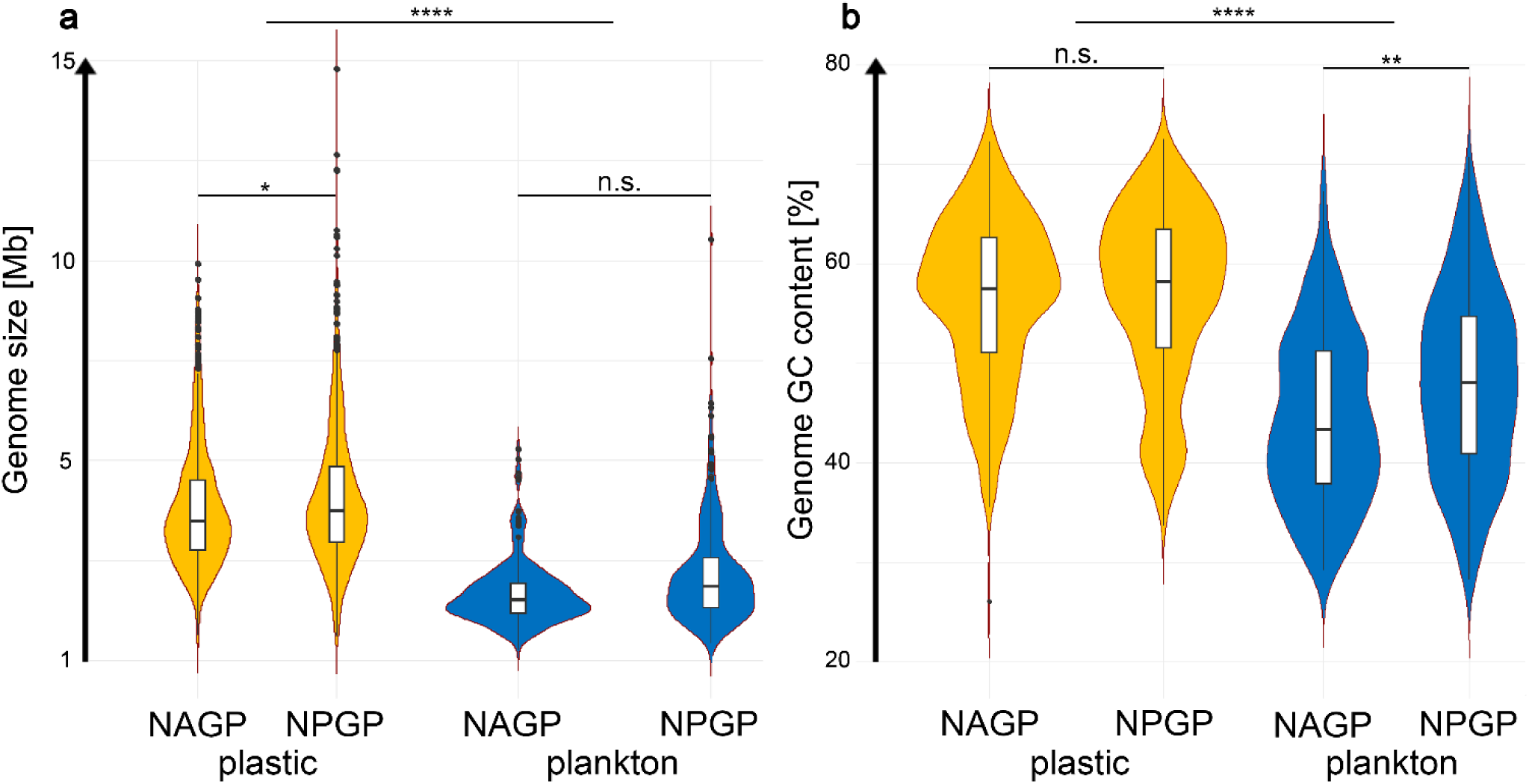
a. Comparison of genome size and **b** GC-content of the plastisphere (n_NAGP_ = 736; n_NPGP_ = 1021) and plankton (n_NAGP_ = 213; n_NPGP_ = 289) at the different oceans. Asterisks indicate adjusted significance levels between the samples according to a pairwise t-test (n.s. not significant, * < 0.1, ** < 0.001, **** < 0.00001).

### Functional comparison of Atlantic and Pacific MAGs

The investigated functional genes were on average 15% more abundant in the plastisphere than in plankton MAGs. Differences in the abundance of (the same) functional genes between plankton and plastisphere were equally distributed in NAGP and NPGP (Fig. 3a). Clustering of the functional fingerprints based on the abundance of all evaluated genes/gene clusters revealed two distinct clusters, which were mostly dominated by classes derived from either the plastisphere or the planktonic metagenomes (Fig. 3b). Exceptions were planktonic *Bacteroidia*, *Actinomycetia* and *Rhodothermia*, which were found next to their relatives in the plastisphere cluster. In the planktonic cluster sporadic *Chlamydiia*, *Gemmatimonadetes*, *Bdellovibrionia*, *Paceibacteria* and single occurrences of *Vampirovibrionia*, *Saccharimonadia*, *Kiritimatiellae* and *Spirochaetia* from the plastisphere were found (Supplementary Table 4).

**Fig. 3.**
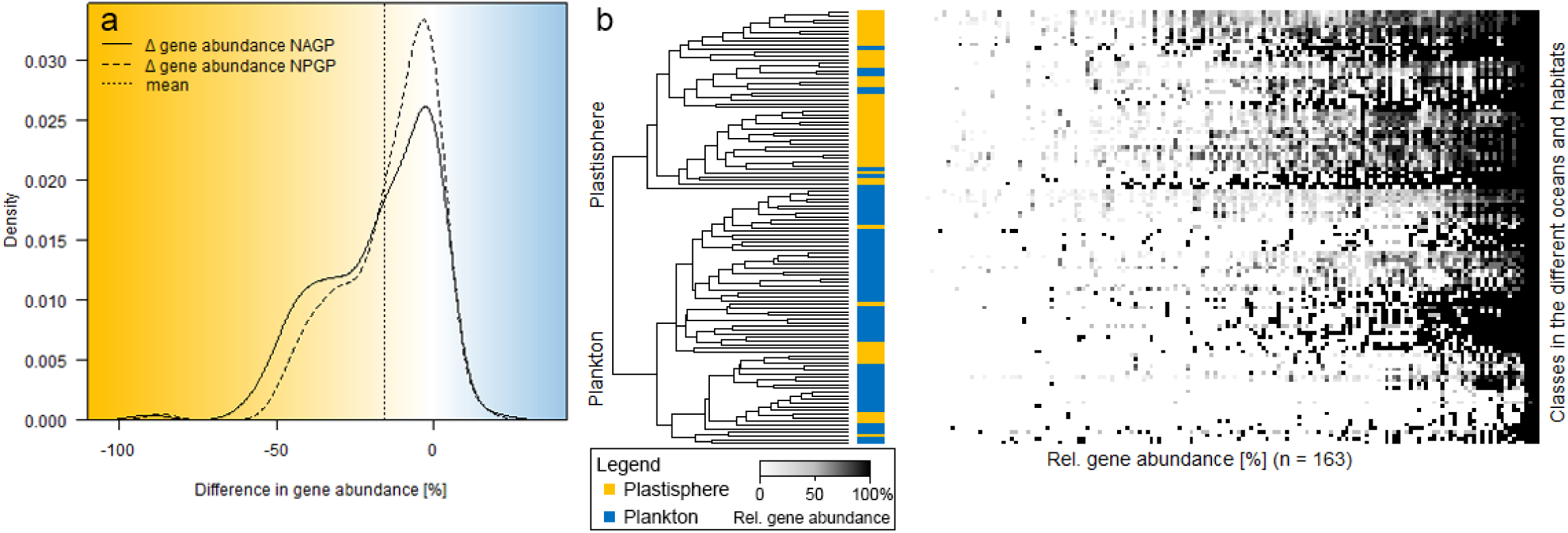
Global functional comparison of relative gene abundance in plankton and plastisphere. **a** Histogram of the difference in relative gene abundance calculated by ***Plankton_g,o_ – Plastisphere_g,o_*** (g - gene; o - ocean), with negative values showing higher gene frequencies in plastisphere than in plankton and *vice versa*. **b** Clustering of the genetic fingerprints (detailed information on the classes and genes are listed in Supplementary Table 4).

From the functional gene fingerprints, we selected 97 genes for a specific analysis of the individual functional potentials, their taxonomic affiliation and their co-occurrence with other functional genes to elucidate how the plastisphere thrives under extreme conditions of the oligotrophic oceans.

### Nitrogen cycling

One of the most pronounced limiting factors in the oligotrophic gyres is bioavailable nitrogen, which is converted into various chemical oxidation states via the nitrogen cycle (Fig. 4a). The abundance of many nitrogen pathways was similar in both oceans and habitats, except for complete assimilatory nitrate reduction (ANR) via nitrate (*NarB*, *NR* or *NasAB*) and nitrite reductase (*NIT-6* or *NirA*) (Fig. 4b). Here 75% of plastisphere MAGs and only 29% of planktonic MAGs had a full ANR potential. On plastic, *Rhodobacteraceae* (23%) and *Phormidesmiaceae* (6%) were the predominant bacterial families capable of full ANR, while in plankton no dominant bacterial families were found. Strikingly, the nitrogen fixation (NF) gene set (*NifDHKENB*) was present in 4 to 7% of plastisphere MAGs compared to less than 1% of planktonic MAGs. The plastisphere MAGs were dominated by *Phormidesmiaceae*, *Nostocaceae*, and *Xenococcaceae* from the class of *Cyanobacteria* (NAGP = 6%; NPGP = 3%) and *Rhodobacteraceae* from the class of *Alphaproteobacteria* (∼ 1%). In plankton, the NF gene set was only present in three MAGs that originate from the NPGP and belong to the classes *Lentisphaeria* and *Phycisphaerae*. All NF-capable MAGs also had a complete ANR pathway. Dissimilatory nitrate reduction (DNR) was an uncommon feature in both habitats, as less than 2% of the MAGs revealed complete DNR. But the DNR gene nitrite reductase *NirBD* was roughly 6 times more abundant in the plastisphere, with *Rhodobacteraceae* (11%) being the major contributor. Genes belonging to the anaerobic process of denitrification were found in up to 7% of the MAGs in both habitats. Only three MAGs from the plastisphere of the NPGP carried the full denitrification potential, whereas the majority had a truncated gene set featuring one metabolic step. The most consistently abundant gene was nitrite reductase *nirK*, which was found in more than 5% of the MAGs of each habitat. In plankton only sporadically, higher abundances of genes were found, such as the nitrate reductase *NarGHI*, nitrous oxide reductase *NosZ* and the nitrification pathway genes *NarGH* and *Hao*.

**Fig. 4.**
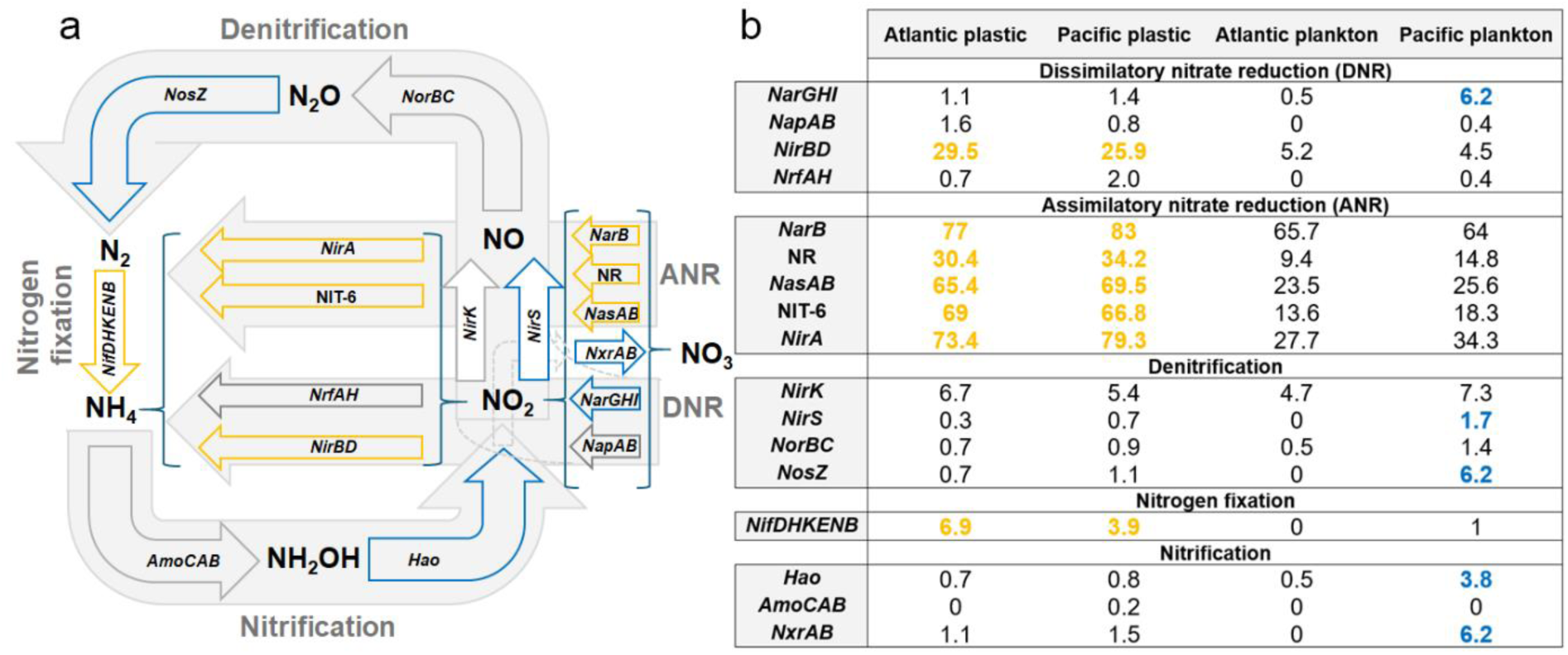
Nitrogen cycle comparison between plastisphere and plankton. **a** Schematic representation of the nitrogen cycle in which the greatest differences between the habitats are highlighted in yellow (plastisphere) and blue (plankton). **b** Abundance table of the respective investigated nitrogen cycle genes. The enzyme names can be found in the Supplementary Table 5.

### Transporters and uptake of essential elements

Individual genes of the cobalt (*cbiMOQ*) and nickel (*nikABCDE*) transport systems were up to 15% more abundant in plastic communities, and genes of the phosphate-specific transport system (*pstABC+pstS*) and alkaline phosphatases (*phoA/B* and *phoD*) up to 42% (Fig. 5a 1-4). With respect to the pst-system 65/63% of the plastic communities of the Atlantic and Pacific possess the whole pst-system, compared to 45/21% of the planktonic communities (Fig. 5a 4). The phosphate-related gene sets further displayed an interesting taxonomic relatedness in the plastisphere, where *Alphaproteobacteria* consistently contained alkaline phosphatases and the pst-system for phosphate uptake, whereas *Bacteroidia* and *Cyanobacteria* typically contained one of the two gene sets. Furthermore, alkaline phosphatases were more commonly found in *Gammaproteobacteria* in plankton, whereas *Alphaproteobacteria* were the most abundant carriers in the plastic communities. In addition, the number of alkaline phosphatase-carrying MAGs in the plankton was rather low, as only 4-5 % of the MAGs in the plankton carried both phosphatases, in contrast to 25-39 % in the plastisphere.

**Fig. 5.**
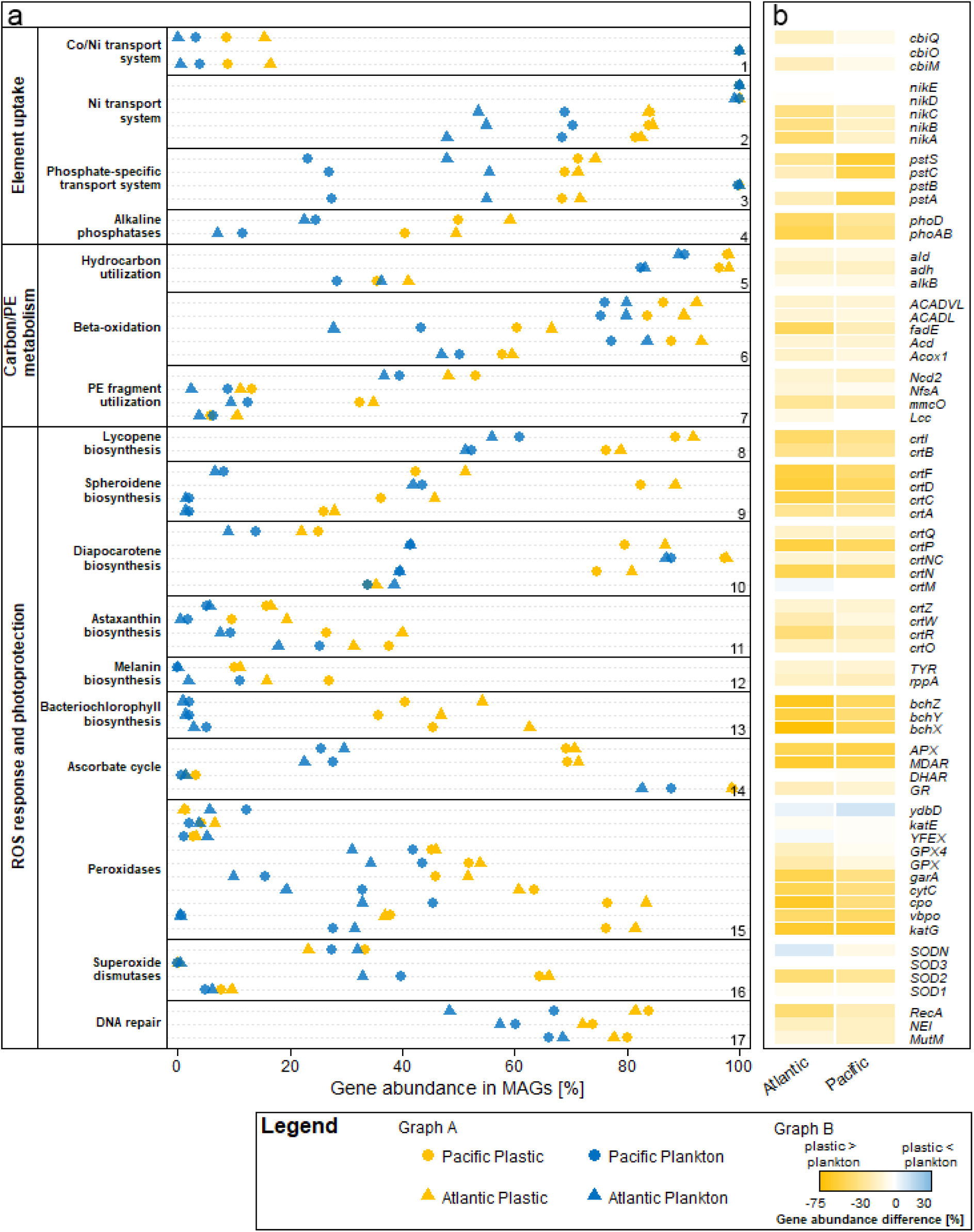
Overview of the abundance of selected functional genes observed in metagenomes from the different oceans and habitats. **a** Relative gene abundance in the MAGs for selected pathways: 1-4) Uptake of relevant essential elements, 5-7) selected Carbon/Polyethylene (PE) utilization pathways and 8-17) ROS response pathways and photoprotection. **b** Gene-resolved heatmap showing the difference in the relative gene abundance calculated by ***Plankton_g,o_ – Plastic_g,o_*** (g - gene; o - ocean) from plot 4a.

### Carbon metabolism and degradation potential

The METABOLIC analysis revealed that the plastisphere bacteria possess a wide carbohydrate metabolism flexibility, in contrast to plankton. Interestingly, number of MAGs carrying genes for complex carbon degradation (cellulase, beta-galactosidase, isoamlyase, hexosaminidase), fermentation and C1 metabolism (glutathione hydrolase/dehydrogenase, formate oxidation, aerobic CO oxidation) were elevated in the plastisphere communities, as well as carbohydrate metabolism genes related to glycolysis, gluconeogenesis, pentose phosphate pathway, Entner-Doudoroff pathway, TCA cycle, glyoxylate cycle and glycogen biosynthesis, amongst others (Supplementary Table 6). Furthermore, the genomic prerequisites for nitrile hydration (nitrile hydratase subunit alpha and beta) were found in 18-20% of the plastisphere MAGs compared to 6-8% of the planktonic MAGs. However, carbon fixation via reverse TCA cycle, hydroxyl-propionate-cycle or the Wood Ljungdahl pathway were not prominent genomic features in either habitat; none of the relevant genes reached more than 10% abundance in the MAGs.

Genes for carbon utilization, such as for alkane monooxygenase (*alkB*), alcohol dehydrogenase (*adh*) and aldehyde dehydrogenase (*ald*), for beta oxidation (*Acox1*, *acd*, *ACADL* and *ACADVL*), RNS acting genes (*NfsA*, *ncd2*) and oxidases were only slightly elevated, with about 4-14% higher occurrences in the plastisphere than plankton (Fig. 5a 5-7). Higher percental differences are found only for individual genes, such as *fadE* (acyl-CoA dehydrogenase), *mmcO* (multicopper oxidase) and PHAases (polyhydroxyalkanoate depolymerases), which are 17-39% more abundant in the plastisphere than in plankton. Interestingly, the taxonomic affiliations of the hydrocarbon utilization and beta-oxidation capable bacteria (only considering the MAGs with the respective full gene set) were quite different between both habitats, while being similar across oceans (Supplementary Table 7).

### Light stress management: changes in pigment patterns

The plastic-associated MAGs displayed a greater genetic potential for phytoene synthase (*crtBI*) with 77/74% of the Atlantic and Pacific MAGs, compared to 40/41% of the planktonic MAGs (Fig. 5a 8). The gene sets for spheroidene (*crtACDF*), diapocarotene (*crtMNNcPQ*) and astaxanthin biosynthesis (*crtORWZ*) were more abundant in plastisphere communities than in plankton (Fig. 5a 9-11). Both enzymes for melanin synthesis were found at elevated levels within the plastisphere, with *TYR* at 11/10% on plastic *vs*. 0% in plankton and *rppA* with 15.8/27% on plastic *vs*. 1.9/11.1% in plankton of the MAGs in the respective Atlantic and Pacific communities. In the plastisphere, we further observed a high abundance of bacteriochlorophyll synthesizing genes (*bchXYZ*), with at least 30% of the MAGs containing the full gene set, but only about 1-2% of MAGs in plankton (Fig. 5a 13). We also found that the combination of bacteriochlorophyll, carotenoid and anoxygenic photosynthetic genes was more prevalent in the plastisphere. In addition, the combination of genes for bacteriochlorophyll and for the photosynthetic reaction center of the anoxygenic photosystem II was linked to the aforementioned ascorbate cycle genes. The respective MAGs primarily belonged to *Rhodobacterales* from the *Alphaproteobacteria* class.

The lycopene pathway was the most abundant among the investigated pigment biosynthesis pathways, allowing for a taxonomic comparison across samples. In plankton, *Poseidonia* primarily possessed the full Lycopene biosynthesis gene set, whereas in the plastisphere *Alphaproteobacteria* dominated. The bacteriochlorophyll-harboring MAGs in the plastic communities were mainly affiliated with *Alphaproteobacteria* (85% in Atlantic and 77% in Pacific plastic MAGs), while the number of planktonic MAGs with this gene set was too low for comparison. In total, only eight planktonic MAGs contained all three genes for bacteriochlorophyll biosynthesis: four gammaproteobacterial, three alphaproteobacterial and one oligoflexial MAG.

### Light stress management: ROS mitigation and DNA repair

The ROS mitigation ascorbate cycle as a whole was not a prominent feature of either plastisphere or planktonic communities (Fig. 5a 14), but the genomic presence of monodehydroascorbate reductase (*MDAR*) and ascorbate peroxidase (*APX*) was approximately two-fold higher in the plastisphere communities, irrespective of the ocean (*MDAR* / *APX* Atlantic plastic 71/70% *vs.* Atlantic plankton 23/30% and Pacific plastic 69/69% *vs.* Pacific plankton 28/26%). Taxonomically, the *MDAR* / *APX* gene set was almost equally abundant in *Alpha*- and *Gammaproteobacteria* in the planktonic MAGs of the Pacific, whereas *Poseidonia* and *Bacteroidia* primarily possessed these genes in the Atlantic. *Alphaproteobacteria* were the most abundant taxonomic group in the plastic MAGs carrying these two genes.

Among the peroxidases, catalase-peroxidase (*katG*), bromide peroxidase (*vbpO*), non-heme peroxidase (*CPO*), cytochrome-c peroxidase (*cytC*), glutathione amide-dependent peroxidase (*garA*), glutathione peroxidase (*gpx*) and phospholipid-hydroperoxide glutathione peroxidase (*GPX4*) displayed higher abundances in the plastisphere. The manganese catalase *ydbD* was the only investigated peroxidase that was more abundant in plankton than in the plastisphere (5.7-12.1% vs. 1-1.3%). In addition, catalase (*katE*) and Porphyrinogen peroxidase (*yfeX*) displayed minor differences in abundance among habitats, and were present only at low levels in all MAGs (<10%) (Fig. 5a 15).

The superoxide dismutases of the Cu-Zn (*SOD1*) and Fe-Mn (*SOD2*) families were more abundant in the plastisphere than plankton (*SOD1* 8-9% in the plastisphere *vs.* 5-6% in plankton and *SOD2* 64-66% in the plastisphere *vs.* 32-39% in plankton). Another superoxide dismutase of the Cu-Zn family (*SOD3*) and nickel superoxide dismutase (*sodN*) were more abundant in the Atlantic plankton than in the Atlantic plastisphere (0.5/32% *SOD3* in plankton *vs.* 0/23% *sodN* in the plastisphere). In contrast, *SOD3* and *sodN* were slightly more abundant in the plastisphere in the Pacific samples (Fig. 5a 16).

UV irradiation and radical species induce DNA damage and can be countered with DNA glycosylases such as *MutM* and *Nei* to correct mismatches, and DNA recombinases, such as *RecA* to fix broken strands. These genes were widely distributed in both environments (48-83% of the MAGs) and were elevated by 9-33% in the plastisphere (Fig. 5a 17).

In summary, a detailed comparison of selected genes revealed that 49 out of 67 genes occurred with at least 10% higher abundance on plastic compared to plankton. These differences were consistent among the oceans and the most striking differences were observed in ROS response and photoprotection and phosphate utilization (Fig. 5b).

## Discussion

We found a high taxonomic similarity between plastic communities in the different oceans. Our taxonomic metagenome analysis revealed that Atlantic and Pacific samples cluster into distinct plastic and plankton groups, which reinforces the existence of a plastisphere^11^, but we could not confirm geographical differences as found in 16S rRNA-based studies^14–16^. However, clustering in this study is based on the frequency of annotated bacterial classes from binned metagenomes, where geographical differences may disappear due to the high level of taxonomic aggregation. The clusters of similar taxonomic classes in the respective habitats indicate a high degree of functionally equivalent species. This aspect is addressed by metagenome binning which allows us to link genes of interest to their respective host genome and thus infer the taxonomic ‘origin’ and the redundancy of the observed functional potential. Following this approach, we were able to show that there is not only a difference in functional potential between plastisphere and plankton as demonstrated by Li, et al. ^13^ but that this is provided by different taxa in the different environments.

In both oceans, we found fewer coding genes, smaller genome sizes and lower GC-content in plankton in comparison to the plastisphere, despite both communities residing in the same environment. Previous studies have shown that the cellular nitrogen costs can be reduced up to 21% in open ocean planktonic microorganisms by decreased GC codon utilization, altered transcriptional regulation and a modified proteome^29,30^. The finding of smaller genome sizes was also shown by Ngugi, et al. ^31^ in ocean plankton and by Michoud, et al. ^32^ in stream biofilms, and both studies linked genomic streamlining to temperature effects. In contrast, we argue that the different niche conditions, e.g. a better nutrient supply, is the driving factor for the selection of species with larger genomes and a higher GC-content, since the plankton and plastisphere species face the same environmental conditions in each respective ocean. This is also underlined by the fact that even within closely related taxonomic families, species living on plastic had a larger genome (Supplementary Table 3).

This hypothesis is also supported by the fact that the plastisphere has a greater potential to control its nutrient supply. The availability of macronutrients such as nitrogen and phosphorus and micronutrients such as cobalt and nickel is a prerequisite for oceanic primary production^33–35^. We found an elevated abundance of nitrogen fixation genes coinciding with increased Co-/Ni transporter genes and alkaline phosphatase and corresponding phosphate-specific transport system genes in the plastisphere. This indicates that the plastisphere has a higher potential to overcome co-limitations by enhanced nutrient uptake and cycling in comparison to plankton. This was postulated from increased abundances of nitrogen-fixing genes^22^ or experimentally shown for alkaline phosphatase activity^36^ and here shown for an extended set of functions related to co-limiting nutrients. An important characteristic of the plastisphere in comparison to plankton is the spatial organization within the biofilm matrix, which improves substrate cycling between organisms but also living in anoxic micro-zones (AMZ) in the biofilm. We found a higher potential for nitrogen fixation in the plastisphere, which is proven to be dependent on AMZ for nitrogenase functioning^37^ and has been measured on planktonic aggregates^38^, sediment particles^39^ or particulate organic material^40^ and is generally stimulated by enhanced surface availability^41^. Thus, the higher potential for nitrogen fixation might be related to micro-zonation within the biofilm. A striking difference of the plastisphere compared with the planktonic metagenomes is the increased potential for nitrate assimilation, a nutrient that is at very low levels in oligotrophic gyres due to vertical stratification^42^. *NirBD*, a gene set involved in dissimilatory nitrate reduction, was also more abundant in the plastisphere, although it can be assumed that these genes are also involved in assimilatory nitrate reduction^43^. However, it remains to be clarified whether these genes are expressed in the plastisphere, as the surface water lacks nitrate and the nitrification potential on plastic was also very low. On the contrary, an increased occurrence of these genes may reflect coastal conditions where these organisms normally thrive, indicating the lack of adaptation needs in the plastisphere in comparison to the plankton environment. In conclusion, our comprehensive evaluation of multiple nutrient-related functions suggests that the simultaneous presence of a higher uptake potential for several limiting nutrients may help to overcome co-limitation, maintain elemental stoichiometry and improve mutualism and nutrient cycling in the plastisphere.

The plastisphere showed potential for more versatile carbon metabolism than plankton, which may reflect the supply of a variety of carbon sources in the biofilm. These might derive from enhanced primary production, as indicated by a high Chl*a* content of the plastisphere^22^ and the release of organic carbon such as chromophoric dissolved organic matter^44^ and polysaccharidic gels^45^, as demonstrated in incubation experiments with microplastics under marine oligotrophic conditions. Thus, higher metabolic diversity in carbon metabolism can indirectly indicate stimulated primary production enabled by nitrogen fixation and a better supply of other co-limiting nutrients.

We found an elevated enzymatic potential for the degradation of complex organic carbon compounds in the plastisphere, which might also be associated with the degradation of plastic polymer fragments. Radicals (ROS, RNS) either from solar irradiation or microbial activity^46^ on the plastic surface potentially induce strand breaks of the polymer-backbone and enhance leaching of compounds from the polymer matrix. This might provide another carbon source for the plastisphere. Leaching studies have shown that plastic can release labile dissolved organic carbon into the ambient seawater during the weathering process, thereby stimulating the activity of heterotrophic microbes^26^. This notion is supported by a comparatively higher abundance of hydrocarbon utilization genes, greater potential for beta-oxidation and presence of genes suggested to be involved in the degradation of PE fragments (laccases, multi copper oxidases)^47^ and other polymer degrading enzymes (PHAases). The microbial utilization of plastic leachates is not well understood, but some taxa are suspected to profit from plastic leachates^48^. Collectively, this supports the notion of metabolic flexibility of the plastisphere suggested in the current study. Such flexibility enables complex interactions among the surface-associated microbial communities on floating plastic particles, hinting towards their autonomy from the surrounding seawater. Furthermore, it illustrates the possibility of utilizing the plastic as source for limiting nutrients. Although, release of plastic constituents might be just a minor source for nutrient supply, perhaps acting as a primer for plastisphere development.

As plastic debris mainly floats at the water surface, microbes in the plastisphere need protection from UV radiation and free radical damage, and related genes were more abundant in the MAGs of the plastisphere than in plankton. Specifically, some ascorbate cycle genes involved in the detoxification of hydrogen peroxide, the majority of peroxidases and DNA repair genes were more abundant in the MAGs of the plastisphere. These findings suggest that the plastisphere is recruited from organisms adapted to a life on plastic debris, floating at the ocean surface for weeks, months or even years while constantly exposed to high light conditions and UV-irradiation.

In the plastisphere, we found bacterial MAGs mainly from the order *Rhodobacterales*, which had the genetic potential to synthesize bacteriochlorophyll and carotenoid pigments, conduct anoxygenic photosynthesis and mitigate radicals through the ascorbate cycle. A cultivation experiment from the same samples from the NPGP plastisphere produced similar pigmented members of the *Rhodobacteraceae* family featuring bacteriochlorophyll and several carotenoids^49^. Our data confirms the higher abundance of the phytoene synthase *crtB* gene in the plastisphere, which was also observed by the previous authors and is now extended to a number of other pigment genes. These findings indicate the presence of a large number of aerobic anoxygenic phototrophs (AAP) in the plastisphere that synthesize ATP by photophosphorylation and have protective mechanisms against ROS. It has been speculated that photophosphorylation is a beneficial adaptation in nutrient-poor marine environments, but it is now assumed that AAPs thrive in productive marine areas or coastal waters^50^. The carotenoids were primarily found in conjunction with RubisCo in *Cyanobacteria*, which indicates their role in photosynthesis in addition to photoprotection. Thus, the plastisphere species seem to benefit from exposure to high light conditions and conserve energy through photoheterotrophy.

In conclusion, our metagenomic analysis supports the hypothesis that the plastisphere represents a new functional unit in oligotrophic oceans. The plastic biofilms from the two oceans share a characteristic taxonomic and functional core composition that is distinct from the surrounding planktonic community. While the planktonic species have streamlined their genomes to cut elemental costs for growth, the plastisphere shows less signs of saving nutrients. On the contrary, the plastisphere has larger, nitrogen-rich genomes that also contain non-essential genes. By virtue of its higher genetic potential to I) actively shape its nutrient supply to overcome co-limitations, II) metabolize a broader range of substrates and III) benefit from high light conditions and engage in photoheterotrophy, the plastisphere can escape the oligotrophic constraints of the gyre and form its own eutrophic niche in an oligotrophic environment. Plastic provides a habitat for non-endemic species, indicating lower evolutionary pressure to adapt to ambient conditions. These species may originate from early colonization in coastal areas, from which plastic mostly originates, or from the enrichment of ubiquitous, underrepresented species from seawater. Our results suggest that the overriding factor for the high functional similarity of the plastisphere microbial communities of the two oceans studied here, is the habitat for biofilm formation. This supports mutualism and nutrient sharing, making the genomic streamlining found in plankton unnecessary. Natural particles are scarce in the studied gyres but play an important role in elemental cycles^41,51^. The expansion of the plastisphere as a result of increasing pollution from plastic particles has the genetic potential to jeopardize the ecological stability of the oceans, in addition to other stress factors such as climate change. This underlines the relevance of ending plastic pollution for the sustainable use of oceans, as indicated in the SDG 14.

## Methods

### Sampling of the plastisphere

Sampling of plastics in the North Pacific garbage patch (NPGP) was conducted on the SO268/3 (MICRO-FATE) transit of the R/V SONNE between Vancouver, Canada and Singapore from May 31, 2019 to July 5, 2019 (Supplementary Table 8). Along this transect, 8 plastic fragments were collected via net sampling and 2.56 cm² of their biofilm was scraped into microcentrifuge tubes using sterile scalpels and frozen in liquid nitrogen.

Sampling of the North Atlantic garbage patch (NAGP) was conducted on the POS536 (DIPLANOAGAP) cruise of the R/V POSEIDON, south west of the Azores from August 26, 2019 to September 4, 2019, and 66 individual small plastic pieces (0,2-1 cm) were collected with a Neuston catamaran (Supplementary Table 8). The whole plastic pieces were stored in RNAlater® and kept refrigerated until further processing.

### Plastisphere DNA extraction and sequencing

DNA of the SO268/3 samples was extracted using the NucleoSpin® Soil Kit (Macherey-Nagel, Germany). SL1 lysis buffer and 150 µL of SX enhancer were used, and sample lysis was performed using a FastPrep® (MP Biomedicals, USA) device at 6.5 m s^-1^ for 35 s in 3 cycles. Between cycles, samples were cooled on ice for 1 min. The following steps were carried out in accordance with the manufacturer’s protocol. Short read sequencing (Illumina) of metagenomic DNA samples was performed by LGC Genomics GmbH (Berlin, Germany) NovaSeq 6000 platform with sequencing depth of approximately 133 M reads per sample resulting in a total of 1 billion reads.

DNA of the POS536 samples was extracted using the AllPrep DNA/RNA Kit (Qiagen, Netherlands). Plastic pieces of an appropriate size were placed in 2 mL BiomedicalsTMLysing Matrix E tubes (MP Biomedicals, USA) with a lysing buffer, containing 2% β-mercaptoethanol and physically disrupted using a FastPrep® (MP Biomedicals, USA), with a single cycle of 30s at a speed of 5500 rpm. The subsequent DNA extraction steps from the lysate was done according to the manufacturer’s guidelines for the AllPrep Kit. Sequencing was performed at the CCGA, Kiel, Germany on an Illumina NovaSeq 6000 platform with a sequencing depth of approximately 16 M reads per sample resulting in a total of 1.1 billion reads.

### Metagenome-assembled Genomes (MAG) - assembly and taxonomic identification in the plastisphere

Assembly and binning were done using the MetaWRAP pipeline (version 1.0.1 for Atlantic samples and version 1.3.2 for Pacific samples)^52^. In short, the metagenomic reads were quality filtered and trimmed using the build in Read_qc module, utilizing Cutadapt^53^ and FastQC^54^. Assembly was done with MegaHIT^55^ and initial binning was done using the three different binning methods available within MetaWRAP (CONCOCT^56^, MaxBin 2.0^57^ and metaBAT2^58^). After the initial binning the bin_refinement module within MetaWRAP was utilized to choose the best version of each MAG from the different previously used binning methods according to a specific minimum completion and contamination. For this study, the thresholds to be included were >50% completeness and <10% redundancy, allowing for a broad bacterial overview of the plastic communities. CheckM^59^ was used for quality assessment of all MAGs and GTDB-Tk version 1.3.0^60^ was used for taxonomic identification.

### Retrieval of planktonic data and comparative metagenomic analysis of plastisphere and planktonic communities

Planktonic metagenomic data was collected within the TARA Oceans Project^61^. The metagenomic raw reads from thirteen TARA Oceans samples nearest to the garbage patches were retrieved from the sequence read archive from NCBI. The TARA Oceans reads were also treated as described in 2.3. This was done to obtain a redundant MAG dataset comparable to the plastic dataset, without losing functional signal between sample types due to dereplication. The planktonic water samples were taken at depths of 5m (9 samples), 40m (two samples), 45m (one sample) and 80m depth (one sample). FastANI^62^ was used to calculate genome-to-genome distance for all MAGs (SI Comp). Open reading frames of each MAG were predicted using Prodigal version 2.6.3^63^. The target genes for this manuscript (Supplementary table 5) were identified in each MAG via hidden Markov model (HMM) profiling using HMMer version 3.1b1^64^ with the HMM search option and a bitscore threshold of 100. For this analysis, multiple gene copies per genome were not considered, but treated as one. In total, approximately 85,000 datapoints were obtained for the investigated 97 genes. The required HMM profiles were sourced from the KOfamKOALA database^65^, which is a customized HMM database of KEGG Orthologs. For the nitrogen metabolism analysis, only whole gene sets within the KEGG reference pathway (https://www.genome.jp/pathway/map00910) were considered, i.e. if all required subunits are present within a single MAG (*NarGHI* -> *NarG*, *NarH* and *NarI*). For the nitrogen fixation specifically, we only included MAGs that contain the complete *NifDHKENB* gene set, thus the catalytic as well as the biosynthetic proteins required for nitrogen fixation^27,66^. High-throughput profiling of all MAGs for functional traits and general metabolic pathways was done using METABOLIC version 4.0^67^, using the METABOLIC-G option.

## Acknowledgements

Mechthild Schmitt-Jansen and Stefan Lips acknowledge funding of the project P-LEACH (https://www.ufz.de/P-LEACH) from the Helmholtz Association, Innovation Pool of the Research Field Earth and Environment in support of the Research Program “Changing Earth – Sustaining our Future” and financial support by the BMBF funded project MICROFATE (GA no.: 03G0268TA & 03G0268TC). Erik Borchert acknowledges financial support from the BMBF funded project PLASTISEA (GA no.: 031B0867A). We acknowledge the logistical support of Annika Jahnke and Mark Lenz as chief scientists of the ocean-going field campaigns SO268/3 and POS536. The authors also acknowledge Peggy Wellner’s (UFZ) assistance in the laboratory and Christoph Rummel’s (UFZ) support with biofilm sampling on board and DNA preparation. We further thank Aaron Beck (Geomar) for critically commenting the manuscript and for language corrections. The metagenomic sequencing of the NAGP samples was done in cooperation with the Competence Centre for Genomic Analysis (CCGA) Kiel.

## Author contributions

SL: design of the work, acquisition of samples and data, analysis and interpretation of data. MSJ: initiation, conception and design of research; acquisition of samples and data, interpretation of data; EB: design of the work, acquisition of samples and data, analysis and interpretation of data. All authors equally analyzed the results and wrote the manuscript.

## Declaration of Interests

The authors declare no conflict of interest.

## Data Availability statement

All metagenomic raw reads and the bacterial plastic MAG assemblies were deposited at the NCBI under the BioProject ID PRJNA901861. The reassembled TARA Oceans MAGs were deposited at the NCBI under the BioProject ID PRJNA1097961.

## Notes

### Competing Interest Statement

The authors have declared no competing interest.

